# Adherent cell remodeling on micropatterns is modulated by Piezo1 channels

**DOI:** 10.1101/2020.11.18.389106

**Authors:** Deekshitha Jetta, Mohammad Reza Bahrani Fard, Frederick Sachs, Katie Munechika, Susan Z. Hua

## Abstract

Adherent cells utilize local environmental cues to make decisions on their growth and movement. We have previously shown that HEK293 cells grown on the fibronectin stripe patterns were elongated. Here we show that Piezo1 function is involved in cell spreading. Inhibiting the Rho-ROCK pathway also reversibly inhibited cell extension indicating that myosin contractility is involved. Piezo1 expressing HEK cells plated on fibronectin stripes elongated, while a knockout of Piezo1 eliminated elongation. Inhibiting Piezo1 conductance using GsMTx4 or Gd^3+^ blocked cell spreading, but the cells grew thin tail-like extensions along the patterns. Images of GFP-tagged Piezo1 showed plaques of Piezo1 moving to the extrusion edges, co-localized with focal adhesions. Surprisingly, in non-spreading cells Piezo1 was located primarily on the nuclear envelope. The growth of thin extrusion tails did not occur in Piezo1 knockout cells suggesting that Piezo1 may have functions besides acting as a cation channel.

## Introduction

Cells detect and respond to mechanical cues with changes in shape, morphology, proliferation, and motility. This process is commonly thought to be mediated via adhesion proteins such as integrins at the adhesion complex (Ingber, 2003; Katsumi et al., 2004; Schwartz, 2010). However, Piezo (1 & 2) ion channels are sensitive to mechanical stress, suggesting that they may serve as a force coupling mechanism at cell adhesions. Piezo1 proteins are the subunits of trimeric non-selective mechanosensitive cation channels (Coste et al., 2010; Coste et al., 2012; Cox et al., 2016; Lewis and Grandl, 2015). Piezo1 is expressed in many cell types where mechanical forces are thought to play a regulatory role (Coste et al., 2010; Syeda et al., 2016). For instance, Piezo1 in endothelial cells can detect shear stress in blood flow enabling alignment along the shear direction (Li et al., 2014; Ranade et al., 2014). Piezo1 is able to transmit information about fluid shear stress to cell nuclei and modulate nuclear size and reorganization (Jetta et al., 2019). Piezo1 channels detect epithelial layer stretching and overcrowding, and can promote division of low density cells (Gudipaty et al., 2017) or extrusion of crowded cells (Eisenhoffer et al., 2012). Piezo1 utilizes local environmental mechanical cues to regulate stem cell differentiation(He et al., 2018). They also respond to osmotic pressure to regulate red cell volume via a Ca^2+^-dependent K^+^ coupling (Cahalan et al., 2015).

Piezo1 protein, originally called Fam38A, was first identified at locations side-by-side with integrins, that suggested Piezo1 and integrins may collaborate to regulate cell adhesion (McHugh et al., 2010). Via Piezo1, substrate stiffness alters the proportion of neurons to astrocytes during neural stem cell differentiation (Pathak et al., 2014). Similarly, in *Xenopus laevis,* substrate stiffness regulates the growth rate and direction of retinal ganglion cells (RGC) (Koser et al., 2016). Activation of Piezo1 increased axon growth, *in vitro* and *in vivo*. On the other hand, a recent study discovered that age-related substrate stiffening caused oligodendrocyte progenitor cells to slow tissue regeneration (Segel et al., 2019), and inhibiting Piezo1 recovered the functional activity of aged cells (Segel et al., 2019).

The micropatterning of adhesive molecules has been widely used to provide mechanical inputs via spatial confinement. Cells grown on narrow stripe patterns stretch themselves along the axis (Suffoletto et al., 2014; Thery et al., 2005). Cells grown on T-shape patterns develop triangular shapes (Suffoletto et al., 2014; Thery et al., 2005). Using FRET based force probes, we previously reported that confinement by patterning the substrates caused a redistribution of tension within actinin during cell spreading (Suffoletto et al., 2014). Cells move faster on narrow patterns than broader substrates (Suffoletto et al., 2018; Vedula et al., 2012). A recent study using Chinese hamster ovary (CHO) cells showed that a Piezo1 mediated Ca^2+^ influx promoted migration of cells that are constrained on the pattern (Hung et al., 2016).

Here we investigated the role of Piezo1 on cell shape and motility on narrow fibronectin stripes. We show that Piezo1 channels are essential for cell spreading on micro-patterns and knockout of Piezo1 eliminates cell expansion. However, inhibiting Piezo1 conductance with drugs leaves the cells with extended thin tail-like features, suggesting the Piezo1 may have additional roles other than mediating Ca^2+^ signaling. Using a cloned Piezo1 expressing HEK cell line, HEK-hP1, we show that Piezo1 protein co-locates with focal adhesion complexes at the tips of spreading cells and this process involves myosin-II contractility via a Rho-ROCK pathway.

## Results

### Piezo1 is required for cell spreading on the micropatterns

We previously reported that HEK cells show elongation along micropatterned fibronectin stripes (Suffoletto et al., 2014). To access the role of Piezo1 in cell shaping, we tested a Piezo1 knockdown in HEK293 cells (P1KO) on the same fibronectin stripes (6 μM wide, 10 μM spacing), and compared the results with control cells and with HEK293T cells stably transfected with EGFP-tagged human Piezo1 (HEK-hP1) (Maneshi et al., 2018). All three cell types attached to the fibronectin surfaces within ~15 min after seeding, and reached a maximum extension in ~2.5 hrs.

Compared with control and HEK-hP1 cells (Fig. 1a, SM movie 1), P1KO cells were not able to stretch to a large extent on the stripes within the experimental period, although they exhibited some ability to move axially (Fig. 1a, SM movies 2). The control cells showed the largest elongation on the stripes (Fig. 1a). HEK-hP1 cells also spread well (Fig. 1a, SM movie 3). The time course of typical P1KO cells linear expansion compared with control cells showed that the control cells expanded for ~3-fold longer than P1KO cells (Fig. 1b). On uniformly coated fibronectin cover slips, the three cell types showed negligible differences in shape (Fig. 1a, lower panel). This suggests that assays based upon open substrates may conceal critical differences in cell physiology.

**Figure 1.**
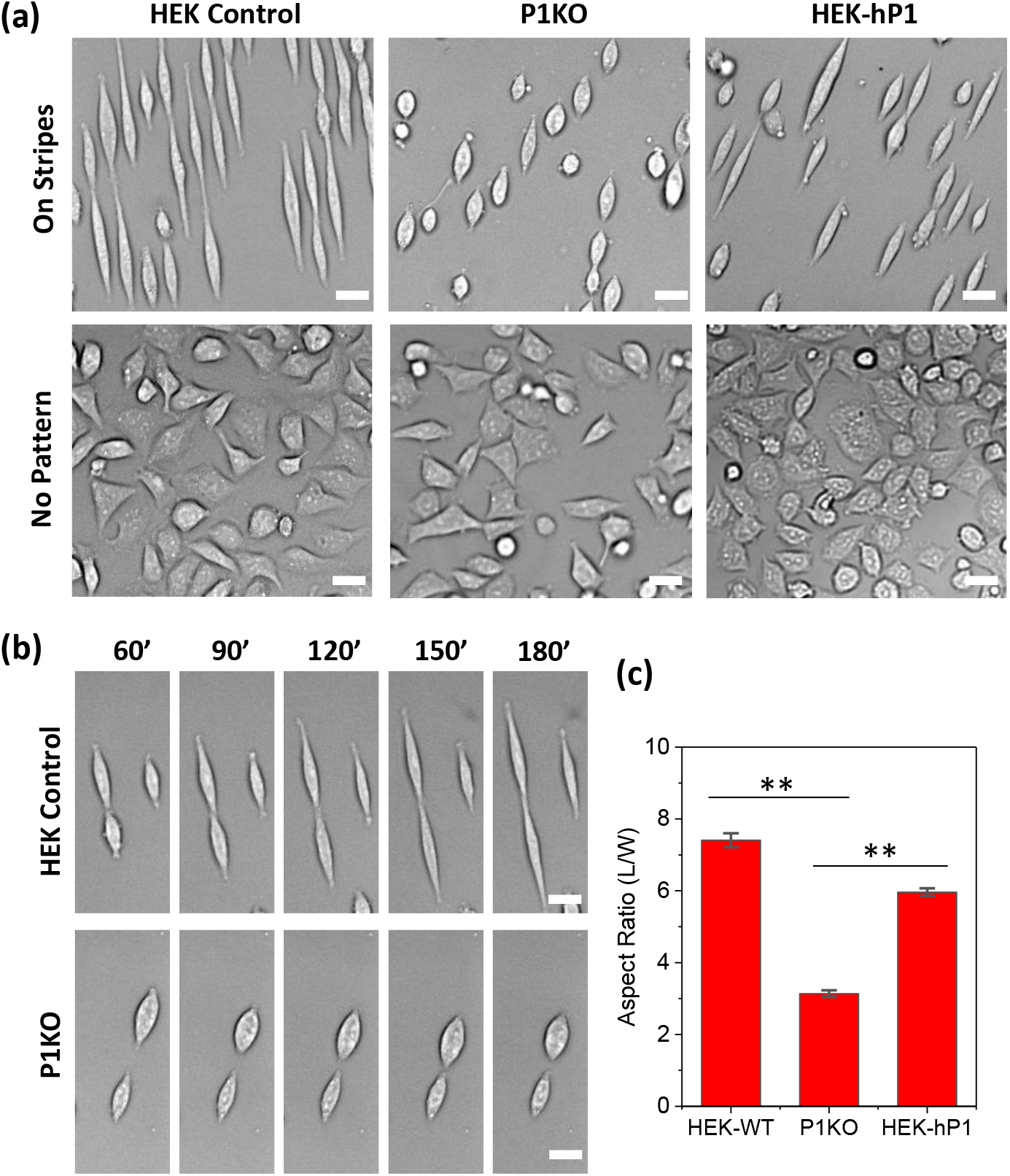
Role of Piezo1 on cell response to micropatterns. **(a):** Images of control, Piezo1 knockout (P1KO), and human Piezo1 expressing (hP1) HEK293T cells following fibronectin microstrips (upper panel), and controls for each cell types without patterning (lower panel). It shows that Piezo1 is required for cells’ response to micropatterns. All images were taken 2.5 hrs after cell seeding. (**b):** Time sequence of cell expansion on stripes of control (upper panel) and P1KO cells (lower panel), showing P1KO cells lost the ability to elongate on the stripes. (**c):** Statistical analysis of the aspect ratio for each cell type (n = 120 from more than 6 experiments for each cell type, \**\*p* < 0.001). Scale bars represent 20 μm.

To quantify the effect of substrate confinement on cell shape, we developed an image processing algorithm to create a binary cell image (see method section for details, SM. Fig. 1). Each cell (the body) was fit with an ellipse, and the ratio of major to the minor axis (L/W) was defined as aspect ratio. The aspect ratios of control and HEK-hP1 cells were significantly larger than P1KO cells (Fig. 1c, n = 120, \*\**p* < 0.001).

### Inhibiting Piezo1 channels inhibited cell spreading

Piezo1 proteins function as mechanosensitive cation channels (MSCs) that open with membrane tension (Bae et al., 2011; Coste et al., 2012; Syeda et al., 2016). Piezo1 mediated Ca^2+^ influx in HEK cells promotes migration (Maneshi et al., 2018). To investigate the functional role of Piezo1 channels on cell elongating, we inhibited the channels with the known Piezo1 inhibitor, GsMTx4 (Bae et al., 2011). Cells did not stretch to a full extent in the presence of GsMTx4 (Fig. 2a). Some cells exhibited an interesting feature; a small cell body connected with long narrow tail-like features (Fig. 2b). While the body length (defined by abrupt change in thickness and width) remained constant, the tails elongated along the stripe (Fig. 2b). They do not expand laterally to cover the patterned surface. As controls, we inhibited mechanosensitive channels using the non-specific inhibitor Gd^3+^, and it too inhibited cell expansion but allowed the narrow tails (Fig. 2a). The tail-like feature was not observed in P1KO cells (Figure 2c, n = 120, \*\**p* < 0.001). These results suggest that Piezo1 may have multiple roles in cell expansion. Cell body expansion requires *opening* of Piezo1 channels with cation currents, likely via a Ca^2+^ influx. However, the growth of thin spikes only requires the *presence* of Piezo1, possibly through their interaction with integrins since they co-localize at cell protrusion edges (see below).

**Figure 2.**
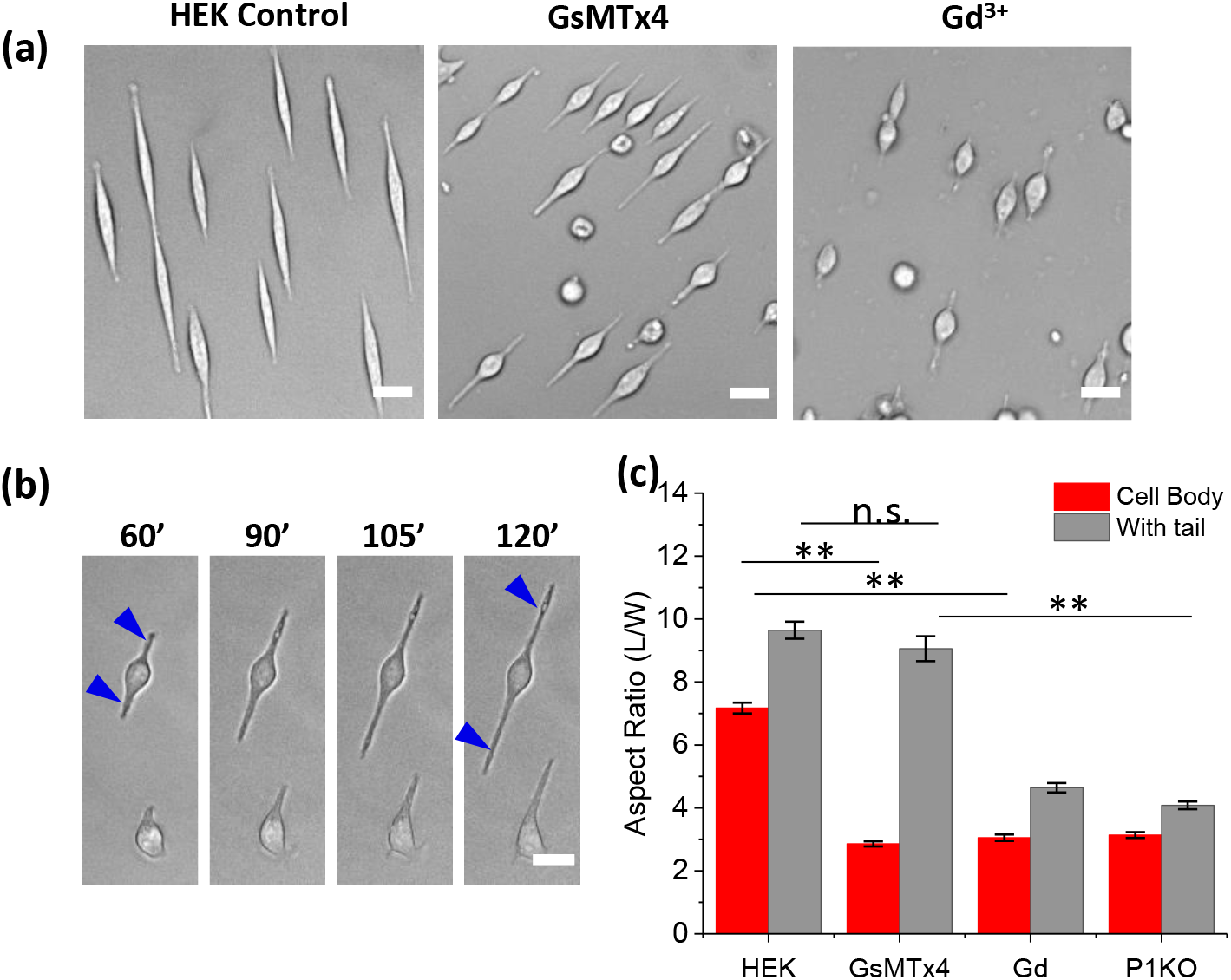
Effect of Piezo1 inhibitors on cell spreading on patterns. **(a)** HEK cells grown on fibronectin stripes in control solution, GsMTx4 (20 μM), and Gd^3+^ (60 μM) for 2.5 hrs, showing inhibiting Piezo1 channels eliminated cell expansion. Drugs were added 15 min after seeding. (**b):** Cell reshaping on stripes in the presence of GsMTx4, showing thin tail-like spikes (indicated by blue arrows) while the cell body remained unchanged. **(c):** Mean aspect ratios (with and without counting the tail length) of cells in media and in the presence of inhibitors (n = 120 for each condition and from multiple experiments, *\*\*p* < 0.001). Scale bars represent 20 μm.

### Piezo1s translocate to the extrusion edges during cell elongating

To investigate the distribution of Piezo1 during cell elongating, we followed GFP labeled Piezo1 in HEK-hP1 cells, and found that Piezo1 plaques continuously translocate from the middle of the cell to the cell extrusion edges during expansion (Fig. 3a, SM Movie 4). When cells reached a maximum length at ~2.5 hrs), the Piezo1 plaques accumulated at the extrusion edges (indicated by red arrows in Fig. 3a). In non-stretching cells, Piezo1 proteins were primarily located on the nuclear envelopes or on the membrane in the middle of the cells (Fig. 3b). Piezo1 density at both edges (Fig. 3a, R1 and R2) was averaged and normalized by the total density over the whole cell (Rtot, Fig. 3a). The time course of the Piezo1 density at the edge and the corresponding body aspect ratio in Fig. 3a is shown in Fig. 3c. An increase in Piezo1 density in extrusion edges consistently coincided with elongation. The mean Piezo1 density at extrusion edges for various cell aspect ratios is shown in Fig. 3d. The edge protein density was consistently higher in stretching than non-stretching cells. In cell treated with GsMTx4, Piezo1 clusters remained in the tails but at slightly lower density (Fig. 3e,f, n = 37, *p* = 0.007). This result suggests that Piezo1 is involved in cell elongation but not via its ion conduction properties. In control cells on a uniform glass surface, Piezo1 is mostly located in the cell body scattered at the cell periphery but we observed no significant number of Piezo1 plaques. The aggregation of Piezo1 into plaques seems to be driven by the physiology required to extend the cells in a confined geometry.

**Figure 3.**
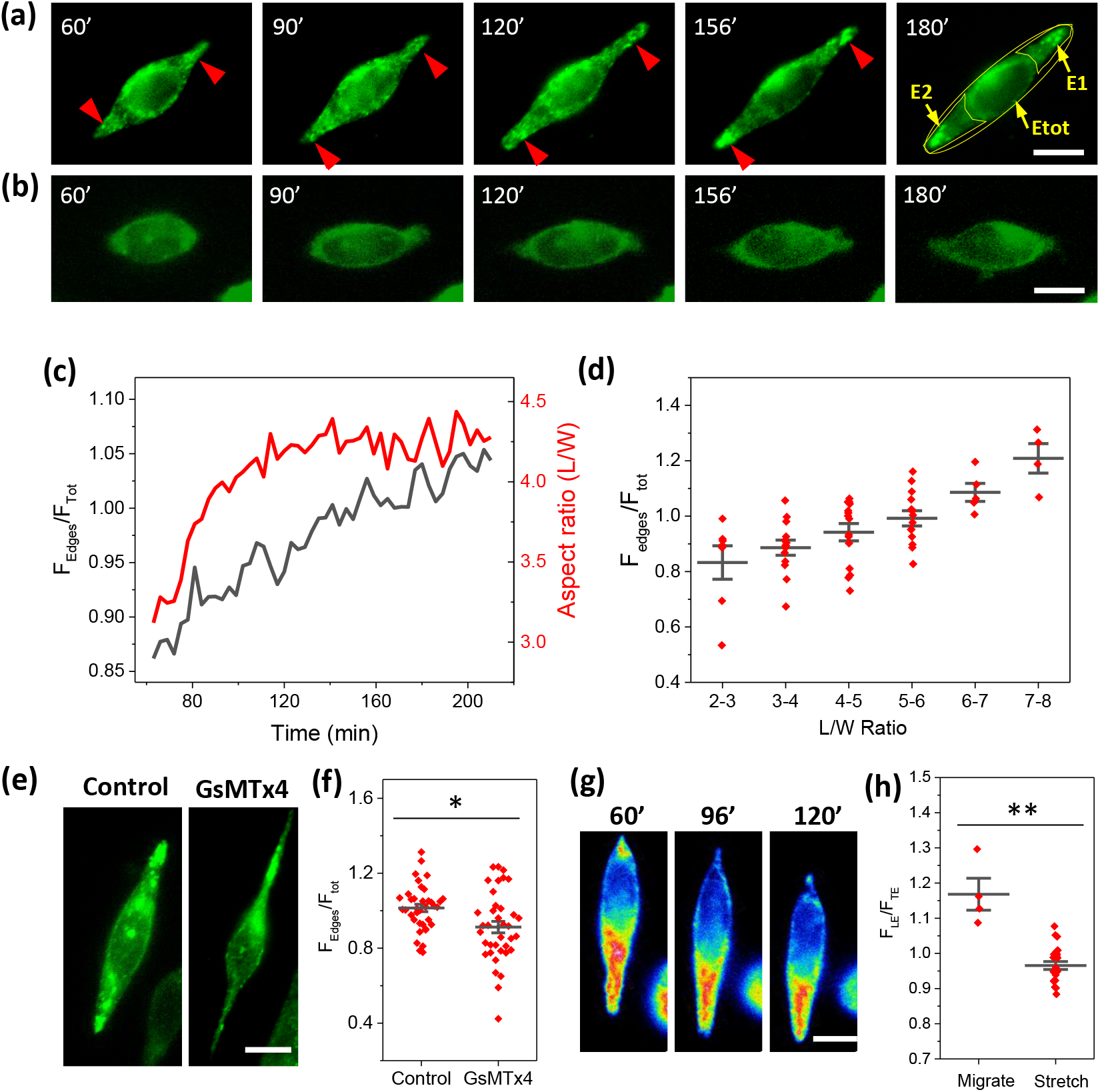
Redistribution of Piezo1 in HEK-hP1 cells on micropatterns. **(a)** Images of typical HEK-hP1 cell extending on the pattern, showing Piezo1 plaques (indicated by red arrows) are present on the extrusion edges of an extending cell. **(b)** shows Piezo1 primarily distributed around the nuclear envelope in non-extending cells. **(c)** Time course of average Piezo1 density over expansion edges (R1 and R2 in Fig. 3a) normalized with whole cell (left axis) is compared with cell aspect ratio (right axis) of the cell in **(a)**, showing increased Piezo1 concentration is correlated with cell expansion. **(d)** Histogram of normalized Piezo1 density at the expansion edges of cells elongated to various aspect ratios, showing higher Piezo1 density corresponding to larger expansion (n = 7, 14, 15, 14, 5, 4, sequentially). **(e)** Fluorescence images show that Piezo1 clusters remained in the tails in the presence of GsMTx4. **(f)** Relative Piezo1 density in the tails following GsMTx4is slightly lower compared with control cells (n = 37, p = 0.007, *\*p* < 0.01). **(g)** Time sequence of Piezo1 distribution in a migrating cell shows a significantly higher Piezo1 density at the leading edge. **(h)** Comparison of Piezo1 density in the leading edge compared to the trailing edge between migrating and stable elongated cells (n = 25 for elongating cells, n = 4 for migrating cell, \*\**p* < 0.001), from multiple experiments. Scale bars represent 10 μm.

Piezo1 density is significantly higher at the leading edge than the trailing edge of migrating cells (Fig. 3g, SM Movie 5). The distribution of Piezo1 in migrating cells compared with immobile but extending cells showed that Piezo1 locates toward the advancing edge of both cell types (Fig. 3h). Cell motility was decreased by GsMTx4 and Gd^3+^, although Piezo1 still concentrated at the advancing cell edges (Fig. 3e). A recent study using Chinese hamster ovary (CHO) cells showed that a Ca^2+^ elevation occurred in migrating cells growing on narrow stripes (Hung et al., 2016). This result is consistent with our earlier report that Piezo1 activity facilitated cell migration (Maneshi et al., 2018).

### Piezo1 co-localized with FAs in extending cells

To see whether the translocation of Piezo1 plaques correlates with the formation of focal adhesions (FAs) during cell expansion, we tested Piezo1 in HEK-hP1 cells co-transfected with mApple-tagged paxillin. Piezo1 plaques at the extrusion edges collocated with FAs in elongating cells but not in resting cells (Fig. 4a). The two labels overlapped in some cells, and appeared side by side in others (Fig. 4b). Although the presence of Piezo1 is necessary for elongation, we cannot tell whether there is an intermolecular reaction between Piezo1 and paxillin or other FA proteins. The FA and Piezo1 plaques only formed at cells extrusion edges in elongated cells. In control cells on glass, Piezo1 is primarily located in the middle of the cell and scattered along the cell periphery. There were no co-localized Piezo1 clusters with mature FAs at the cell edges (Fig. 4c). Piezo1 can be recruited to FAs(Yao et al., 2020), our results show that co-localization of Piezo1 and FAs does not occur when the cells are in the relaxed state.

**Figure 4.**
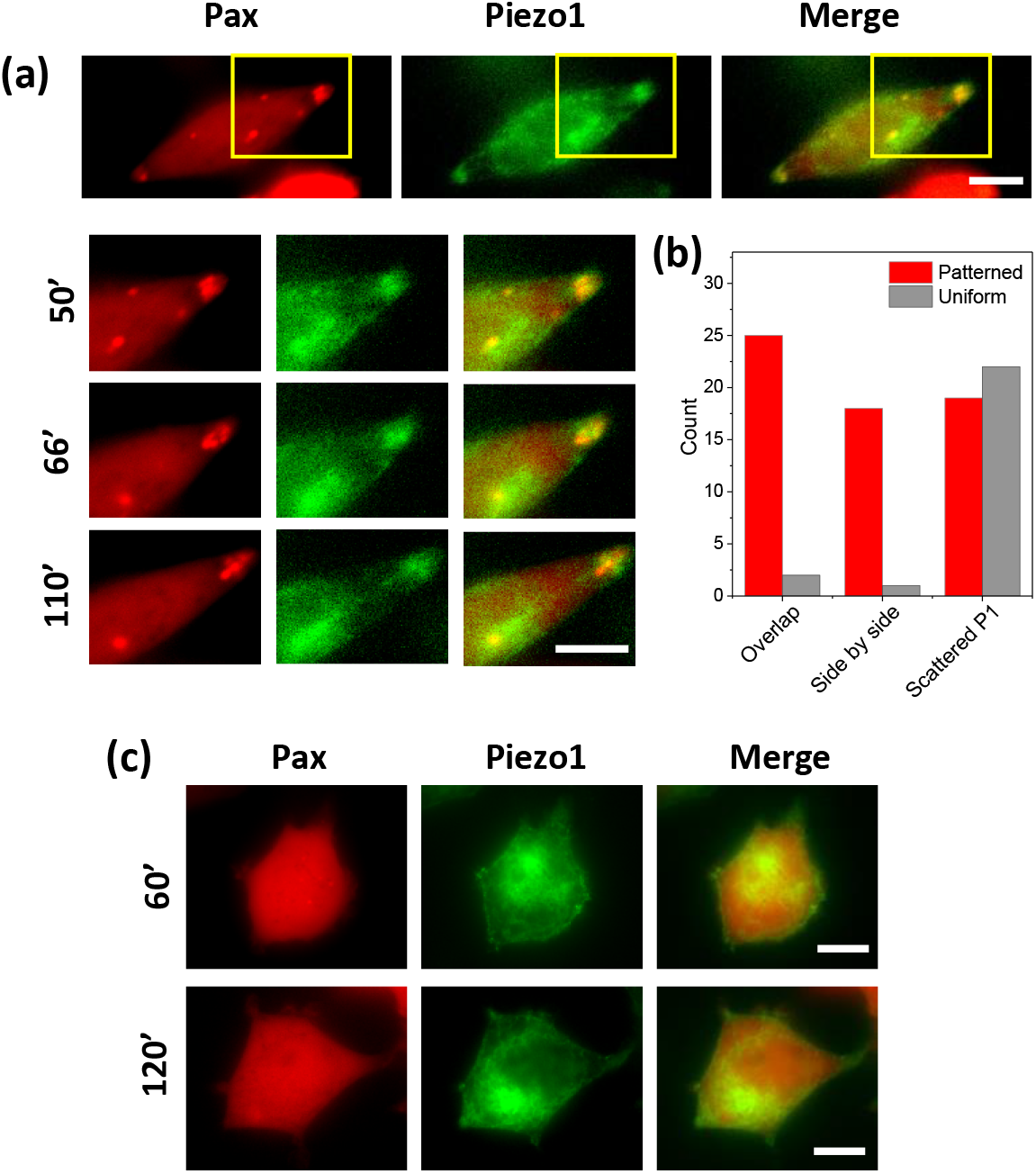
Piezo1 plaques correlate with FAs at the cell extrusion edges on the pattern. **(a)** Images of a typical HEK-hP1 cell (Piezo1, green) co-expressing mApple-paxillin (red); the zoomed view shows the localization between Piezo1 plaques and FAs at the extrusion edge at various times during cell elongation. **(b)** Statistical data shows that the two proteins collocated in most elongated cells, but side by side in some cells. **(c)** Images of paxillin and Piezo1 in a control cell (without pattern) shows that there are no mature FAs present at the cell edges. Scale bars represent 10 μm.

### Rho-ROCK pathway is involved in cell spreading

We previously reported that change in cell shape and expansion over patterned surfaces is a force sensitive process that requires functional actomyosin Rho-ROCK pathways (Suffoletto et al., 2014). We tested whether Rho-regulated myosin-II contraction force plays a role in translocation of Piezo1 proteins and the opening of Piezo1 channels. We inhibited Rho-ROCK pathways with the ROCK inhibitor Y27632, and it reversibly inhibited cell expansion (Fig. 5a). In the presence of inhibitor, the cell body shrank, but the tail-like feature remained. After washout, the cells regained full expansion (Fig. 5a). The effect of inhibitor was consistent in HEK-hP1 and control cells (Fig. 5b, also see SM Fig. 2). This suggests that Rho-II associated contractile forces affect Piezo1 channel functions required for cell full spreading. The distribution of Piezo1 in the cell edges was not affected by Y27632, despite cell shrinkage (Fig. 5c), cells exhibited similar tail-like feature as they had under GsMTx4 (Fig. 3e). The translocation of Piezo1 to the extrusion edge of the cell was not reversible. This result is consistent with our previous report that Rho-II associated contractility could lead to changes in membrane tension that opens Piezo1 channels causing cell expansion.

**Figure 5.**
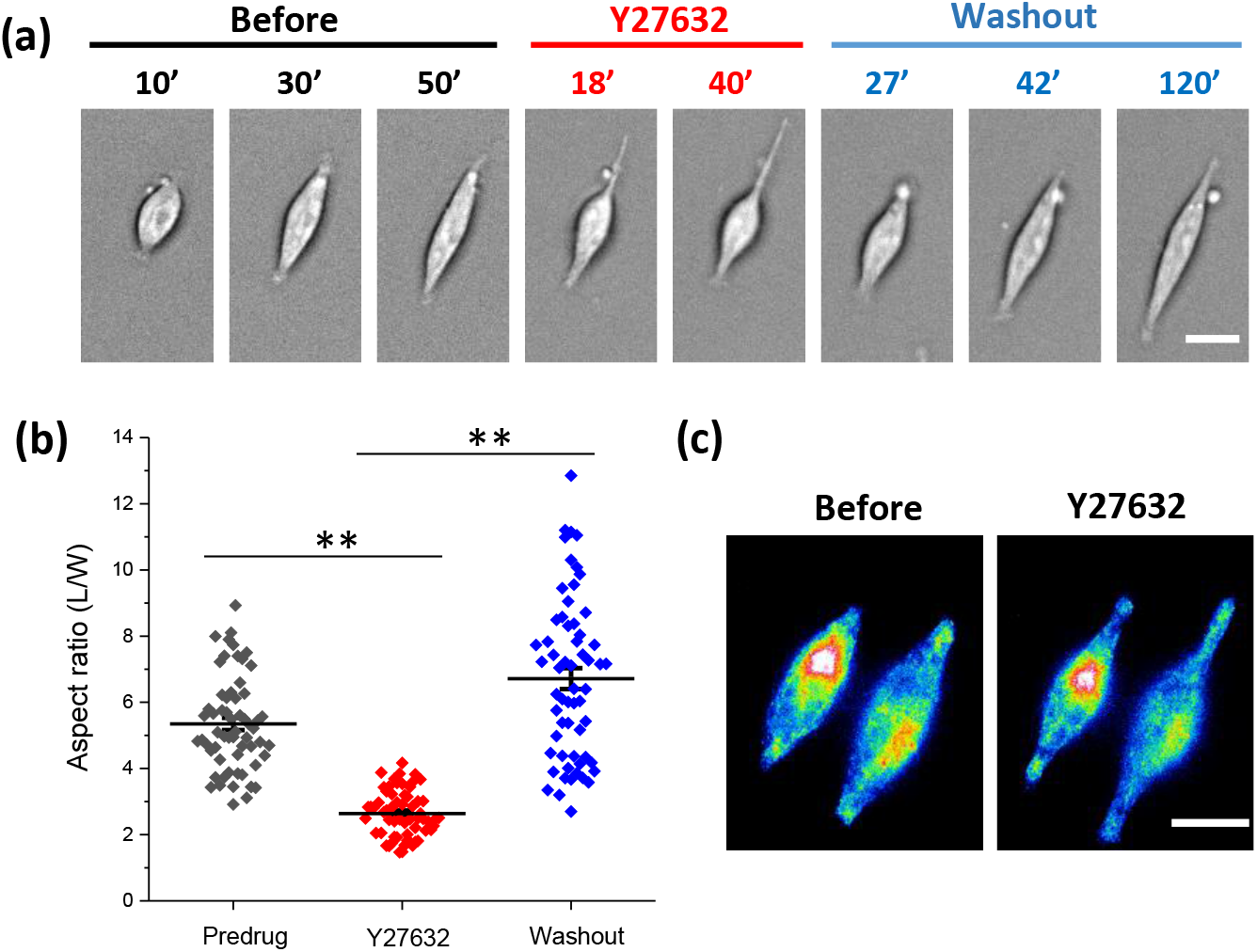
Effect of Rho-ROCK inhibitor on cell elongation. **(a)** Cell’s response to Rho-ROCK inhibitor Y27632 followed by washout, showing Y27632 reversibly inhibited cell elongation. The times measured from the beginning of each process, seeding, drug application, and washout, respectively. Scale bar represents 10 μm. **(b)** Mean aspect ratio for pre-drug treatment, 20 min after treatment, and 100 min after washout, shows that the drug effect is statistically significant (n = 60 for each condition from 6 experiments, \**\*p* < 0.001). **(c)** GFP images of a typical HEK-hP1 cell before and after application of Y27632, showing Piezo1 proteins remained at the cell edges although the cell body contracted under the drug. Scale bars represent 10 μm.

## Discussion

Adherent cells respond to mechanical cues such as spatial confinement on patterned substrates. Our results show that knockout of Piezo1 inhibited cells’ ability to respond to the pattern (Fig. 1), demonstrating that Piezo1 is involved in this process. Blocking Piezo1 channels with inhibitors, GsMTx4 and Gd^3+^, inhibited the overall cell expansion measured by aspect ratio of cell body on the pattern, but cells grew a thin, long, tail-like feature. This result that inhibition of Piezo conductance is not the same as the knockout indicates that Piezo1 may have roles beside ion conductance on cell shape. The full spreading of cells requires Piezo1 mediated current. During cell expansion, Piezo1 clusters moved from the cell soma to the extrusion edges. Piezo1 plaques co-localized with the FA complex at the extrusion edges elongating cells. The body expansion, but not the thin tails, was eliminated by the ROCK inhibitor, suggesting that both Piezo1 and Rho-ROCK signaling are functional in fully spreading cells.

Piezo1 is a mechanosensitive ion channel, reputed to regulate cell physiologies via Ca^2+^ signaling(Coste et al., 2010; Coste et al., 2012). Wild-type HEK293T cells express endogenous Piezo1 channels (Cox et al., 2016), that could be opened during cell expansion activating Ca^2+^-dependent downstream pathways leading to cell remodeling. Inhibiting Piezo1 channel activity inhibited cell expansion, indicating the ion influx mediated by Piezo1 plays a role consistent with the observation that fluid shear stress caused a Ca^2+^ influx in HEK293T cells through Piezo1 channels(Maneshi et al., 2017; Maneshi et al., 2018). However, inhibiting Piezo1 conductance only inhibited the full cell spreading; cells still grew the long thin spikes (Fig. 2). Piezo1 may have roles in cell elongating in addition to its functions on Ca^2+^ influx. Piezo1 protein, originally called Fam38A, was first identified at locations side-by-side with integrins, suggesting that Piezo1 and integrins collaborate to regulate adhesion(McHugh et al., 2010). As Coste et al(Coste et al., 2010) pointed out, disrupting integrin signaling does not affect Piezo1 current suggesting that activation of Piezo1 could lead to integrin activation, not other way round (Coste et al., 2010). Our results demonstrate that thin cell extrusion growth does not require the opening of the channels.

During cell spreading, Piezo1 clusters continuously translocate from soma to extrusion edges forming plaques at the cell tips (Fig. 3). The relative protein density at the spreading edges in stretching cells is higher than non-stretching cells. In migrating cells, the Piezo1 distribution is polarized and mainly located at the leading edge. The redistribution of Piezo1 proteins was only observed in the cells on narrow stripes that imposed a mean tension gradient on the cell. On uniform substrates, Piezo1 is mostly located in the middle of the cell. A previous study by the Rosenblatt group shows that Piezo1 overexpression occurred at the cell edges of sparse regions where the cell membrane was expanded (Gudipaty et al., 2017). The redistribution of Piezo1 proteins is sensitive to the stresses in cells such as cytoskeletal stress and correlated membrane tension.

It is widely recognized that cells use transmembrane integrins at focal adhesions to translate extracellular matrix (ECM) mechanical cues to the cytoskeleton to facilitate cell remodeling (Ingber, 2003; Katsumi et al., 2004; Schwartz, 2010). Our study shows that Piezo1 plaques co-localize with mature FAs at cell expansions. Although Piezo1 proteins exist in the thin tail-like features, no mature FAs were observed in those domains. Without the patterns, neither distinct Piezo1 plaques nor mature FAs were observed along the cell periphery (Fig. 4). A recent study has shown that Piezo1 is recruited to FAs in normal cells but not in transformed cells including HEK(Yao et al., 2020), that observation was done on a uniform substrate, consistent with our control cells.

Our results further show that cell expansion on the pattern can be reversibly inhibited with the Rho-ROCK inhibitor Y27632, although the tail-like features remained (Fig. 5). This suggests that Rho-ROCK activated Myosin-II contractile force affects Piezo1 mediated Ca^2+^ influx required for full cell expansion but not Piezo1 translocation. It is known that Myosin-II contractile force regulates the formation of focal adhesions and actin assembly through a Ca^2+^-dependent MLC kinase (MLCK)(Eddy et al., 2000). It has been reported that traction forces alone can activate Piezo1 to generate local Ca^2+^ flickers in neural stem cells(Ellefsen et al., 2019). This process involves Light Chain Kinase regulated myosin-II(Ellefsen et al., 2019). Thus, Piezo1 mediates Ca^2+^ influx, that in turn can also promote Myosin-II contractile forces and maturation of FAs (Ye et al., 2014). With Piezo1 inhibitors, there are nascent FAs since the lamellipodial actin extrusion and formation of nascent FAs does not involve a significant contraction force (Giannone et al., 2007).

In conclusion, Piezo1 channels function to translate mechanical cues that affect cell spreading. A change of local tension due to cell growth on the confined surface area triggers Piezo1 migration from the soma to the extrusion edges. Piezo1 has at least two roles at the cell edge. They open to create a Ca^2+^ influx, that activates Ca^2+^-dependent contraction forces in the Rho-ROCK pathway, enabling FA formation and cell expansion. Inhibition of Piezo1 influx does not block the translocation of proteins, nor the growth of thin tail-like features showing that inhibition of ion influx not the same as the knockout, so that Piezo may have other enzymatic activities.

## Materials and Methods

### Substrate patterning

Parallel fibronectin stripes 6 μm wide with 10 μm spacing were printed on a glass coverslip using standard microprinting techniques. Briefly, a PDMS stamp was fabricated using soft lithography. Coverslips were cleaned in a boiling mixture solution of ammonium hydroxide and hydrogen peroxide for 10 min, and were rinsed with ethanol and dried under UV light. The coverslips were then treated with plasma for 3 min to increase hydrophilicity. To print fibronectin on the coverslip, fibronectin (50 μg/ml) was applied over the stamp and left for 1 hr, and rinsed with PBS. The stamp was then pressed in contact with the coverslip. To reduce non-selective cell attachment, the patterned cover glass was immersed into a blocking reagent (0.2 % Pluronics F-127) (Sigma-Alrich) for 1 hr. Finally, a ring-shaped thick PDMS wall was bound to the coverslip to form a dish containing a patterned glass bottom. The patterns were examined with NHS-Fluorescein dye (ThermoFisher Scientific) using a fluorescent microscope.

### Cell culture and Piezo1 cell lines

Human embryonic kidney (HEK293T) cells (ATCC), and HEK cells stably expressing EGFP-tagged human Piezo1 (HEK293T-hP1) (Cox et al., 2016), and Piezo1 knockout (P1KO) (kind gift from the Patapoutian group) were cultured in Dulbecco’s Modified Eagle Medium (DMEM) complemented with 10% fetal bovine serum (FBS) and 1% penicillin and streptomycin in culture flasks. For visualizing FAs, HEK-hP1cells were co-transfected with 0.2 μg plasmid DNA of paxillin-mApple (generous gift from Michael Davidson) using transfection reagent Effectene (Qiagen, Valencia, CA), and were cultured for 24 hours.

The cells in the flask were trypsinized and seeded onto the patterned substrates. Unattached cells were gently washed away using PBS after 10 min. During experiments, the chip was placed in a stage-top incubator (INUB-ZILCSD-F1-LU, Tokai Hit Co., Ltd, Japan) maintained at 37 °C and 5% CO_2_. B/W imaging was done in DMEM without phenol red DMEM (Gibco, TX) to avoid background fluorescence. Isotonic saline solution was used for imaging GsMTx4 treated cells since DMEM reduces the potency of the peptide (Bae et al., 2011).

### Fluorescence imaging

The images were acquired using an inverted microscope (Axiovert 200M, Zeiss) with a CCD camera (AxioCam, Zeiss). Fluorescence images were obtained with two filter sets: GFP filter set (Ex: 365/40; Em: 445/50 nm) and RFP filter set (Ex=550/25, Em=605/70), and a 63x oil immersion objective. The B/W images were obtained using a 20x objective.

### Cell morphology evaluation and statistical analysis

The change in morphology of the cell was quantified by its length to width aspect ratio, calculated using ImageJ (NIH) with the plugin MATLAB. Brightfield images were processed with several erosions and dilations in MATLAB and the regions of interest (cell bodies) were identified using the ‘adaptive threshold’ algorithm built in Image Segmenter app. The output image was then thresholded to yield a binary image (SM, Fig. 1b). The number of erosions and dilations remained the same for all the images in order to remove bias from processing. This procedure does not include the ‘tail-like’ structures and rendered only the full expansion of the cell body. The total length with tails was separately measured. The binary images were then transferred to ImageJ for fitting each cell body with an ellipse (SM, Fig. 1c,d). The ratio of the major axis to the minor axis (L/W) of an ellipse was defined as the aspect ratio of a cell.

### Statistical analysis

The aspect ratios were averaged over multiple cells in each panel and from multiple experiments. A minimum of six experiments were performed for each condition. Data are shown as the mean ± standard error of the mean (s.e.m.). Statistical analysis was performed using the two sample t-test (The drug effect was calculated using paired sample t-test). Values of *p* < 0.001 were considered statistically significant.

### Solution and chemicals

GsMTx4 was purified and diluted in saline to a final concentration of 20 μM. Gadolinium chloride and Y27632 (all from Sigma-Aldrich, St. Louis, MO) were prepared to final concentrations of 60 μM and 30 μM, respectively.

## Author Contributions

D.J. and M.R.B.F. designed and performed the experiments and analyzed data; F.S. interpreted data and edited manuscript; K.M. analyzed data; S.Z.H. designed the experiments, analyzed and interpreted data, and wrote the manuscript.

## Acknowledgements

This work was supported by National Science Foundation (CMMI-2015964 and CMMI-1537239). We thank Dr. Philip Gottlieb at University at Buffalo for helpful discussions.

