## Supplemental Figures for "Adherent cell remodeling on micropatterns is modulated by Piezo1 channels"

### Supplemental Materials

#### Supplemental figures

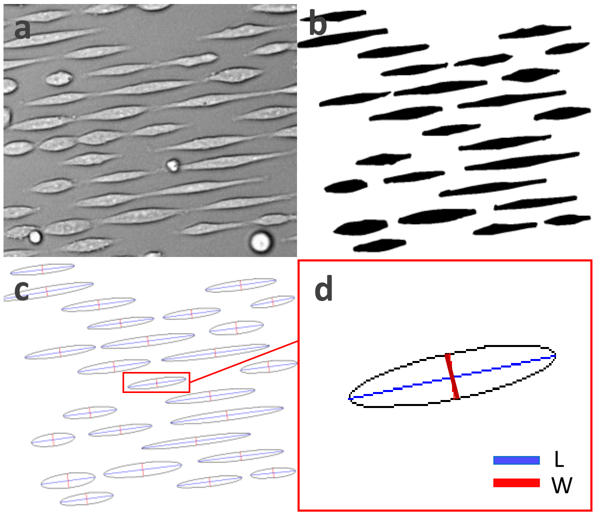


**SM Figure 1. Image processing steps for quantifying cell morphology. (a, b)** Brightfield images were transformed to a binary image using ImageJ with plugin MATLAB and the region of interest (cell body) was identified using the 'adaptive threshold'. **(c, d)** Individual cells were fitted with ellipses and the ratio of major and minor axes of the ellipses defined as the aspect ratio.

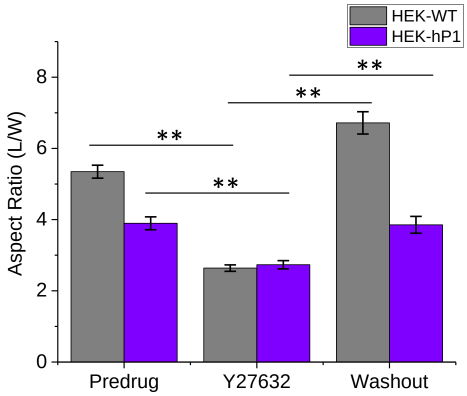


**SM Figure 2. Effect of Rho-ROCK inhibitor on HEK-hP1 cells.** Mean aspect ratio for pre-drug treatment, 20 min after treatment, and 60 min after washout for HEK-hP1 cells (n = 26, $^{**}p < 0.001$) and control HEK cells (n = 60, $^{**}p < 0.001$). It shows the effect of inhibitor was consistent in both HEK-hP1 and control cells.
